# Dissecting Light Sensing and Metabolic Pathways on the Millimeter Scale in High-Altitude Modern Stromatolites

**DOI:** 10.1101/2022.02.10.479979

**Authors:** Daniel Gonzalo Alonso-Reyes, Silvina Galván, José Matías Irazoqui, Ariel Amadio, Diogo Tschoeke, Fabiano Thompson, Virginia Helena Albarracín, María Eugenia Farias

## Abstract

Modern non-lithifying stromatolites (STs) on the shore of the volcanic lake Socompa in the Puna are affected by several extreme conditions. Although STs were proposed as ecologic models for understanding stress response and resilience in microbial ecosystems constituting a window into the past, our knowledge of ST function is still nascent. The present study assesses for the first time light utilization and functional metabolic stratification of STs on a millimeter scale through shotgun metagenomics. In addition, a scanning-electron-microscopy approach was used to explore the community. Our results demonstrated that Bacteroidetes and Cyanobacteria play major roles as ST builders and primary producers to sustain a diverse community of heterotrophs. STs manifest a high occurrence of genes for the synthesis of UV-protecting pigments, the cryptochrome-photolyase family (CPF), and rhodopsins in the surface layers. Three different ecologic niches involving the use of light in energy production were defined. Calvin-Benson and Wood-Ljungdahl pathways were proposed as the main mechanisms for carbon fixation. Several genes account for the microelectrode chemical data and pigment measurements performed in previous publications. We also provide here an explanation for the vertical microbial mobility within the ST described previously. Finally, our study points to STs as ideal modern analogues of ancient STs.

## INTRODUCTION

Microbial stromatolites (STs) are stratified organosedimentary structures built as a result of the trapping, binding, and/or *in-situ* precipitation of minerals linked to the metabolic activities of microorganisms (Walter, 1976; Burne and Moore, 1987). These structures are considered the earliest complex ecosystems on Earth and geological records suggest biogenic evidence dating from about 3.5 billion years ago (Schopf and Packer, 1987; Schopf, 2006; Nutman et al., 2016). STs were ubiquitous until the beginning of the Phanerozoic, after which time their abundance rapidly declined as a result of the incipient grazing and burrowing activities of metazoans and protistans (Walter and Heys, 1985; Awramik and Sprinkle, 1999). Though present analogs of STs are scarce, molecular-genetic analyses have begun to uncover the complexity for some specimens—including from settings in the Bahamas (Khodadad and Foster, 2012; Casaburi et al., 2016); in Shark Bay, Australia (Wong et al., 2015, 2018); and in Yellowstone Park, USA (Pepe-Ranney et al., 2010).

More recently, non-lithifying modern STs were discovered on the shore of the remote volcanic lake Socompa at 3,570 m above sea level in the Puna (Argentine Andes), which region is affected by extreme environmental conditions. Accordingly, investigators propose these STs as the best proxy analogs of the ancient forms (Farías et al., 2011, 2013). Indeed, the Puna region is an arid to semiarid environment withstanding the most elevated doses of global solar radiation on the planet (Piacentini et al., 2003; NASA, 2014). The monthly average for the daily irradiation values reaches 6.6 KWhm^−2^ per day, a level that is among the highest in the world (Duffie and Beckman, 2013). The Puna values are clearly above the radiation levels detected on the Tibetan plateau (29.° N, 91.1° E; 3,648 m), with a monthly mean ultraviolet (UV) index of over 16 in July (Ren et al., 1999). In addition to the resulting UV-B chemical stress (Ruggieri et al., 2010), the lake water is alkaline (pH 9) at salt concentrations of about 10% (w/v); plus the arsenic content of the STs reaches 18.5 mg.L^−1^ and that of the water 32 mg.L^−1^ (Farías et al., 2013). Favorable conditions, however, are provided by a nearby hydrothermal stream that supplies nutrients and maintains the water-column temperature at a relatively stable level (around 26 °C). The Socompa microbial ST systems could thus be considered as an ecological model for understanding stress response and resilience in microbial ecosystems as well as a wide window to be opened into the past.

The mineral fraction of STs has been previously characterized along with bulk-layered microbial diversity of those structures by means of targeted culturing and the cloning and sequencing of 16S-rRNA-primed PCR products (Farías et al., 2013; Toneatti et al., 2017). Likewise, pigment and microsensor analyses have been performed through the various layers (50 mm-deep), revealing steep vertical gradients of light, oxygen, hydrogen sulfide, pH, and pigments in the porewater. Three different zones were defined: an initial UV-stressed zone between 0 and 2 mm (the two oxic layers); a second transitional oxic zone between 2 and 5 mm dominated by infrared (IR) light (layers 3 and 4), and a third anoxic low-IR and sulfide-rich zone between 5 and 7 mm (layers 5 and 6). Nevertheless, the functional metagenomics repertoire of the expected pathways rich in ST metabolism in each of these zones is still unknown.

Microbial interactions and biochemical cycling have been proposed to occur on a fine, millimeter scale (Kirk Harris et al., 2013), with steep nutrient gradients supporting niche differentiation, thus enabling different microorganisms within specialized niches to interrelate with each other and the environment (Macalady et al., 2008; GRÜNKE et al., 2011). Previous work has led to the proposal that some putative niches (Toneatti et al., 2017)—for example, phototrophic consortia near the surface layer of the mat—are the main source of energy production fuelling the anoxic layers. In fact, a common feature of the microbial mats of the high-altitude Andean lakes (HAALs) is the low abundance of Cyanobacteria (Albarracín et al., 2015), which paucity raises the question as to which other group(s) of microbes acts as a primary producer. Phototrophic microorganisms transduce light into energy by two general systems: chlorophyll-related pigments and retinal-based microbial rhodopsins (Pinhassi et al., 2016). The primary production in microbial mats was largely attributed to chlorophyll and/or bacteriochlorophyll photosynthesis, whereas the role of the microbial rhodopsin in mats—though more widely studied in marine ecosystems (Gómez-Consarnau et al., 2019)—is uncertain. Nonetheless, the autotrophic repertoire of Socompa STs is largely unknown.

Recent work has begun to clarify the fundamental role of photoreception in the HAAL microbial mats: indeed, the microbialites from Socompa and Diamante lakes had the higher abundances of photolyases than those from other environments, and those abundances were correlated with higher UV-B–radiation intensities (Alonso-Reyes et al., 2020). Kurth et al. (2021) demonstrated that rhodopsins could be a fine alternative source of energy for the La Brava and Tebenquiche mats above the low participation of cyanobacteria in light conversion. Albarracín et al. (2016b) reported the isolation and characterization of the first rhodopsin from a Socompa ST, proposing that the use of rhodopsin to counteract stressful conditions could be a strategy of those communities situated at the top layer, where light is fully available but UV stress is maximal. More intriguing were the observations that a fully phototactic layer exists in the mat, capable of moving vertically through the ST in response to changes of radiation intensities (Farías et al., 2013).

The analysis of separate stromatolite layers by shotgun metagenomics offers a higher resolution in delineating the functional capability of putative niches to be discovered. Of particular interest is the poor presence of bacterial nitrifiers and the lack of corresponding nitrifying genes (Farías et al., 2013), suggesting incomplete or replaced modes of nitrogen cycling in Socompa stromatolites. Thus, metagenomic analyses at higher resolution would be required to ascertain how communities overcome potential bottlenecks in these systems. Accordingly, the aim of the study reported here was to assess for the first time light-utilization profiles and functional metabolic stratification within STs on a millimeter scale by means of metagenomics.

## 1. MATERIALS AND METHODS

### 1.1. Sample Collection

The samples analyzed in this study were collected at noon in February of 2011 (Southern-Hemisphere summer) in parallel to the samples analyzed by Farías et al. (2013). Permission for sample collection was granted by the Ministerio de Ambiente y Desarrollo Sustentable, Salta, Argentina (number 000388; 17–09– 2010).

Columnar round-dome–shaped stromatolites are found on the southern shore of Socompa Lake (Fig. 1, Panel A), where a stream of hydrothermal water enters the lake. Unlike in the winter, when the STs are covered because of deep snowfalls, during the summer the higher evaporation rates expose the top 10–30 cm to dry air and direct sunlight (Fig. 1, panels B, C). The top six layers of the sample, extending from the surface down to a depth of 7 mm, were dissected with a sterile scalpel based on their defferent individual coloration (Fig. 1, panels D, E): the top layer (0.3 mm in depth) was white with little cracks and pinkish patches; the second layer (at 0.3–1.5 mm depths) was dark-green; the color of the subsequent layers, whose thickness varied with depth, changed between light and dark brownish.

**Fig. 1.**
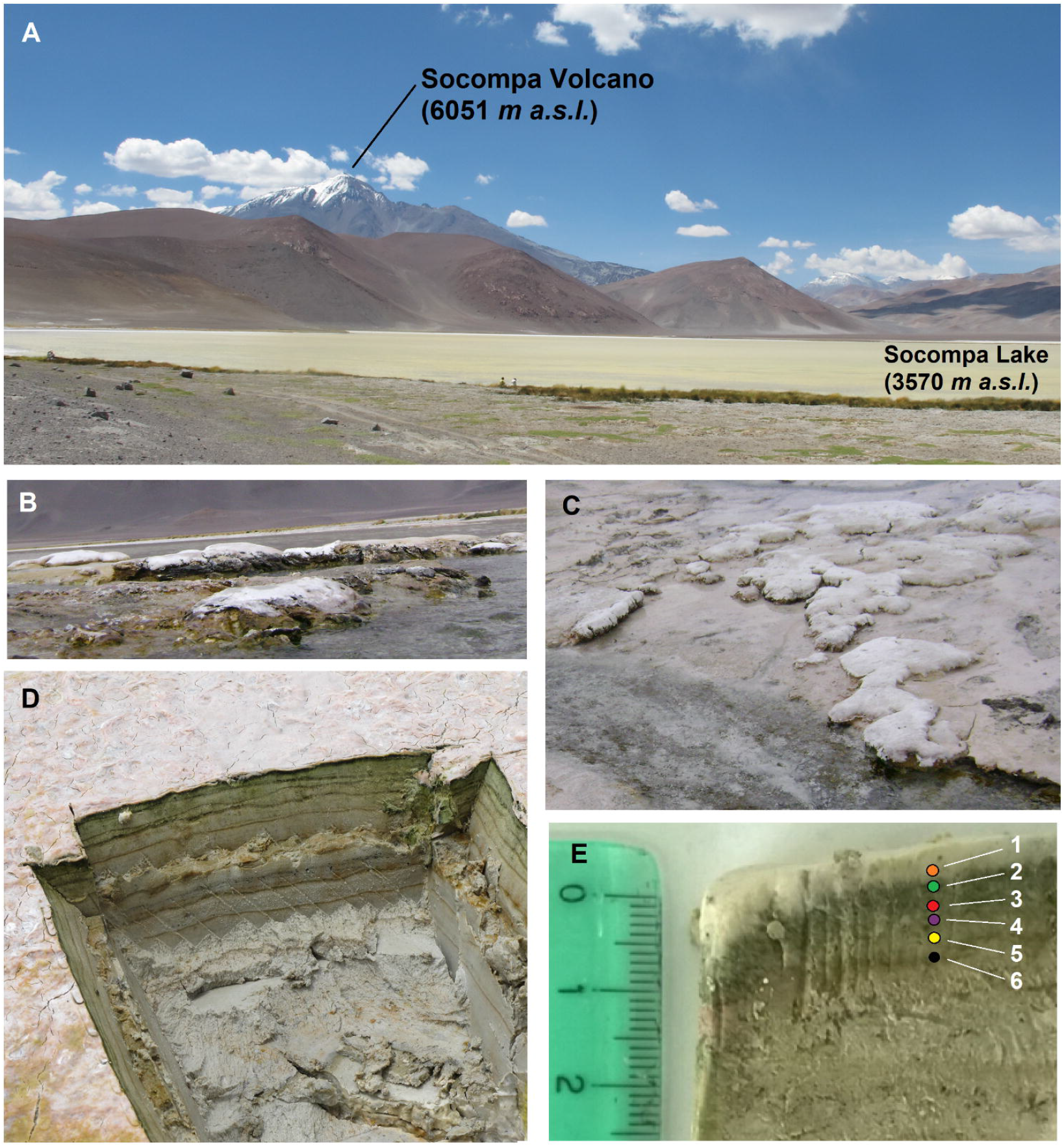
Environmental setting of Socompa Stromatolites. Panels A–C: Sampling site at the shore of the alkaline Socompa Lake containing stromatolites of domical and columnar morphology. Panel D: Transverse cut of the sample with clearly visible laminations. Panel E): Magnification of stratification revealing a pinkishwhite layer on the surface (1, orange dot), a dark-green layer (2, green dot), a light-brownish layer (3, red dot), a dark-brownish layer (4, purple dot), a light-brownish layer (5, yellow dot), and a dark-brown layer (6, black dot) in successive strata below. m.a.s.l., meters above sea level

Samples for DNA analysis were frozen in liquid nitrogen, stored in the dark, and processed within a week of sampling. The DNA of each layer was extracted with the Power BiofilmTM DNA Isolation Kit (MO BIO Laboratories, Inc.) along with a bead-tube method and the inhibitor-removal technology R according to the manufacturer’s instructions.

### 2.2. Sequencing, quality control, and downstream analyses

Metagenomic libraries (n = 6) were prepared through the use of the Nextera XT DNA Sample Preparation Kit, according to the manufacturer’s instructions. The DNA of those libraries was purified with AMPure XP beads and quantified by the fluorimetric Qubit dsDNA High Sensitivity Assay system (Life Technologies). The quantification of the libraries was performed with the 7500 Real Time PCR (Applied Biosystems, Foster City, CA, USA) and the KAPA Library Quantification Kit (Kapa Biosystems, Wilmington, MA, USA). The library-size distribution was accessed with the 2100 Bioanalyzer (Agilent). The six metagenomes (one per layer) were sequenced by Illumina MiSeq paired-end sequencing (2 x 150 base pairs) at the Laboratory of Microbiology (Federal University of Rio the Janeiro). Reads from all layers were quality-trimmed with Trimmomatic v. 0.36 (Bolger et al., 2014), and then a coassembly of all samples was obtained by means of SPAdes v. 3.11.1 (Bankevich et al., 2012) with the “--meta” option. The coverage of each contig in each layer was obtained through the use of Bowtie2 v. 2.2.4 (Langmead and Salzberg, 2012) with default parameters and SAMtools v. 1.6 (Li et al., 2009). The genes of over 500 bp were predicted in all the contigs by Prodigal v. 2.6.3 (Hyatt et al., 2010) with the “--meta” parameter.

The functional annotation of genes was performed by HMMER3 v.1.b2 (Eddy, 2011), with the profiles obtained from pfam (https://pfam.xfam.org/) and made in-house through the hmmbuild option of HMMER3 (Table S1D), from representative sequences aligned with the MAFFT online aligner (/https://mafft.cbrc.jp/alignment/server/). For the LOV domain, we used the sequences and coordinates indicated in the alignment of Crosson et al. (2003). HMMER3 hits and/or counts were filtered by an e-value of 0.001 and 70% of coverage. The resultant sequences were blasted against the NR-NCBI database for confirmation of their function and those with conflicted annotation were discarded and not considered as counts. Sequences annotated as rhodopsins were further classified through the use of the MicRhoDE online database for microbial rhodopsins (Boeuf et al., 2015). In contrast, sequences identified as photolyases were classified in subfamilies through the sequence-similarity-network method (Gerlt et al., 2015). The gene abundances were normalized for all layers in the following way:

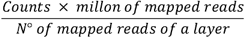

The gene abundances were plotted as a heat maps by means of the the Pheatmap v. 3.6.3 R package (Kolde, 2019). The characterization of the architecture of multimodular proteins was performed by querying the sequences against the pfam database and results plotted through an in-house R script.

To find the layer-by-layer diversity, the reads were queried against the nr-NCBI protein database with DIAMOND (Buchfink et al., 2014) and the results subsequently processed by MEGAN6 v.19.1 (Huson et al., 2016). The genes of the functional analysis were searched through BLAST against the online National-Center-for-Biotechnology-Information database for taxonomical classification.

### 2.3. Scanning-electron-microscopy Analysis

The morphology of the round, dome-shaped STs was analyzed by scanning electron microscopy (SEM), whereas the elemental composition was evaluated by energy-dispersive X-ray spectroscopy. To that end, samples were fixed with Karnovsky’s fixative (2.66% [v/v] paraformaldehyde and 1.66% [v/v] glutaraldehyde) in sodium phosphate buffer (0.1 M, pH 7.2) overnight at 4 °C. Thereafter, the fixed samples were washed three times with phosphate buffer and CaCl_2_ and fixed with osmium tetroxide (2% [w/v]) overnight. After fixation, the samples were dehydrated with aqueous ethanol (30% [v/v]) followed by absolute acetone and critical-point dried (Denton Vacuum model DCP-1). The specimens were sputtered with gold and then observed under vacuum with a Zeiss Supra 55VP (Carl Zeiss NTS GmbH, Germany) scanning electron microscope.

## 2. RESULTS AND DISCUSSION

### 2.1. Sequencing, trimming, assembly

The Illumina-MiSeq paired-end sequencing of the six metagenome samples generated 19,005,054 raw reads, ranging from 1,079,347 raw reads (Layer 1) to 2,090,410 raw reads (Layer 2). Table S1A summarizes the trimming results. The assembly resulted in 33,607 contigs over 500 bp, but was highly fragmented, with only 667 contigs over 10 Kbp. The number of reads mapped to the assembly in each layer was also low. The second layer had the highest percentage of mapped reads (50.1%), while the sixth layer had the lowest (21.9%).

### 3.2. Biodiversity throughout the layers as assessed by metagenomics and microscopical methods

The microbial diversity was first determined in the bulk ST and then layer by layer by means of 16S-rRNA–gene-amplicon metagenomics (Farías et al., 2011, 2013; Toneatti et al., 2017). In this work, we applied shotgun-metagenomic sequencing with an aim at obtaining a more detailed analysis of the taxonomic and functional diversity within the ST layers (Fig. 2). We also related our taxonomic results to data from scanning electron microscopy (SEM) and SEM coupled with an energy dispersive X-ray detector (Fig. 3 and Supplementary Files S1–S4).

**Fig. 2.**
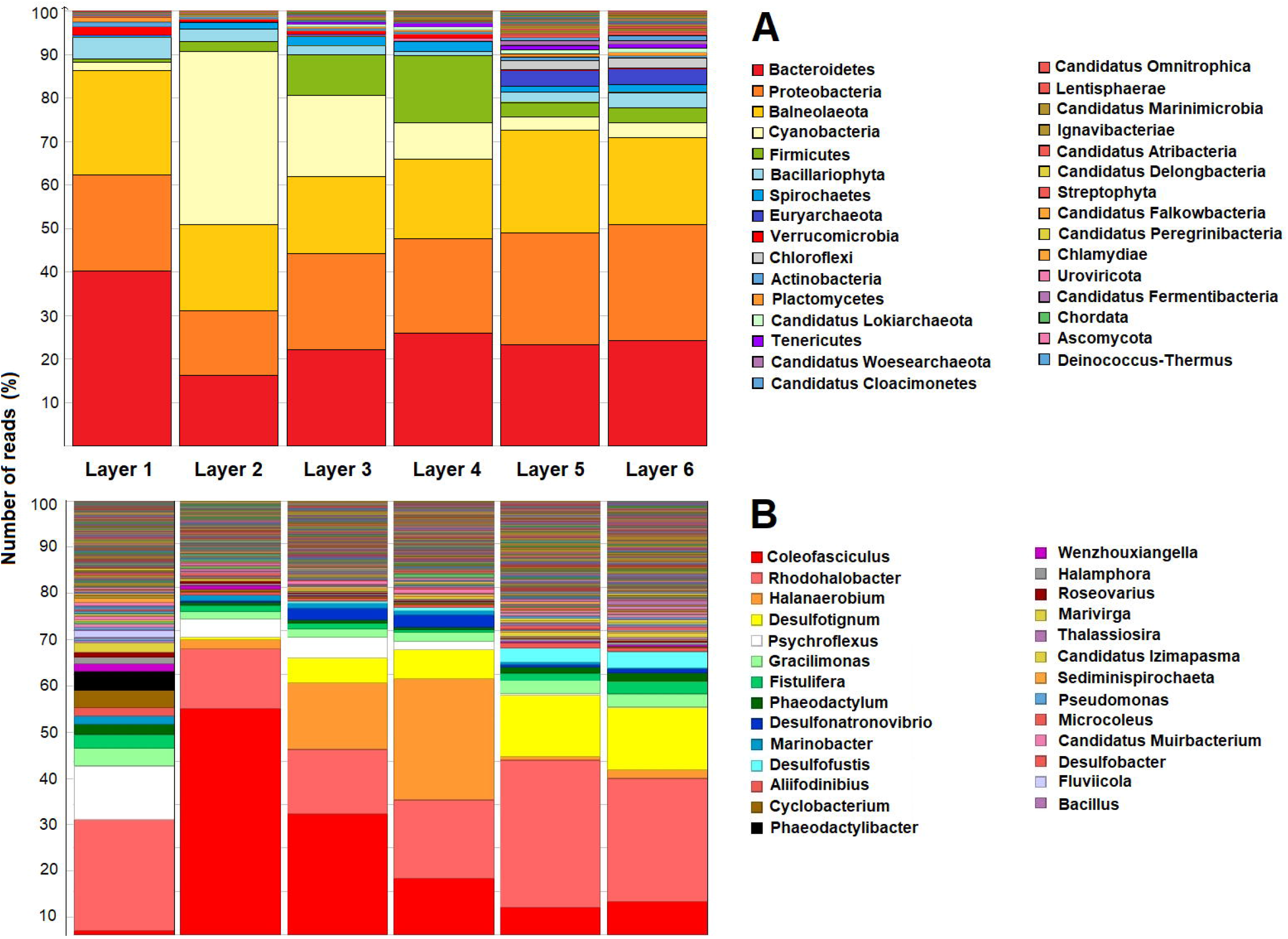
Stacked bar chart indicating the taxonomical profile layer by layer. In the figure, for each layer indicated above (Panel B) or below (Panel A) the vertical bars, on the right *ordinate* is plotted the number of reads for each microbial taxonomic assignment within all six layers, expressed as a percentage of the relative abundances obtained from Illumina shotgun reads, generated through MEGAN 6, and classified against the NCBI-NR database. The fraction of a given bar occupied by each color reflects the percent of that Phylum or Phyla, (Panel A) or Genus (Panel B) within the total consortium according to the color code to the right.

**Fig. 3.**
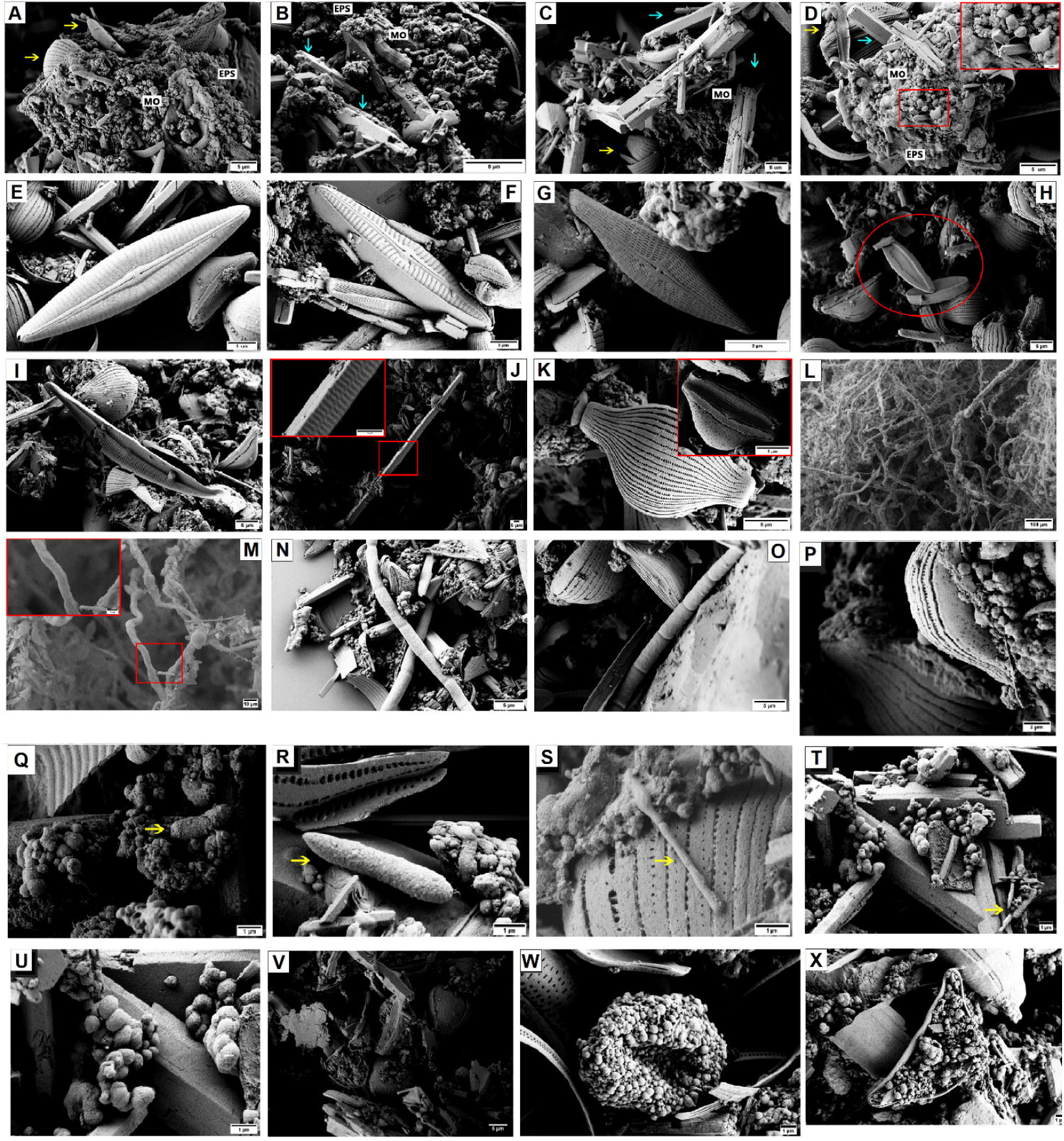
SEM micrographs of the various layers of the stromatolite. Panels A–D: polyform aggregates present in different zones of the stromatolite. Panels A and B: Microbial aggregates near the mat surface: microorganisms are embedded in extracellular polymeric substances (EPS) associated with diatom frustules (yellow arrows) and aragonite crystals (cyan arrows) confirmed by X-ray diffraction. Panels C and D: Microbial aggregates from deeper areas; a higher concentration of aragonite crystals (cyan arrows) is observed, corresponding to the deepest layers, as well as diatoms (yellow arrows). The image with high magnification (Panel D, red box) reveals the microbial interaction (bacilli) with inorganic matter (crystals) in detail owing to the presence of the EPS. Panels E–K: Colonization of diatoms: Panel E: *Naviculaperotii* (length range, 47.2 μm; width range, 10.1 μm). Panel F: *Naviculaperotii* (length range, 41.7 μm; width range, 9.3 μm). Panel G: *Naviculaveneta* (length range, 20.0 μm; width range, 4.8 μm). Panel H: *Naviculahodgeana* (length range: 15.2–16.8 μm; width range, 3.46–5.40 μm). Panel I: *Amphora copulata* (length range, 48.9 μm; width range, 6.5 μm). Panel J: *Synedra* sp.(length range, 48.9 μm; width range, 6.5 μm). Panel K: *Halamphora* sp. (length range, 21.7 μm; width range, 12.7 μm). Panels L–O: Colonization of Cyanobacteria. Panels L–M: *Coleofasciculus* sp. *Microcoleus* sp. Panels N–O: Red boxes in panels J, K, and M indicate amplified images of *Coleofasciculus* filament. Panels P–U: Suggestive heterotrophic morphotypes. Panels P–R: prokaryotes found in the oxic zone. Panels D–F: Prokaryotes found in the anoxic zone. Panel B: *Rhodohalobacter* sp. (yellow arrow). Panel S: *Psychroflexus* sp. (yellow arrow). Panel T: *Gracilimonas* sp. (yellow arrow). Panel U: *Halanaerobium* sp. (yellow arrow). Panels P–R: Deinococcus-Thermus coccoid cells, found in both areas (oxic and anoxic). Scale bar: 2 μm (Panel P), and 1 μm (panels Q–R). Diatom frustules colonized by heterotrophic bacteria of variable morphologies (panels V–X)

The dominant heterotrophs of the surface layer were mainly Bacteroidetes, Balneolaeota, and Proteobacteria. The bacteroidete *Psychroflexus* sp. (Bowman et al., 1998), a psychrophile, was second in abundance (Fig. 2, Panel B) understandably owing to the low temperatures experienced by the ST surface overnight. The abundance of chlorophyll and bacteriochlorophyll phototrophs was low, with many of those members belonging to the eukaryotic phyla Bacillariophyta (diatoms). Most of those diatoms belonged to the order Naviculales. Accordingly, SEM images visualized microbial aggregates associated with diatoms and carbonate minerals in all the layers. Bacteria of variable morphologies (mainly cocci and bacilli) were observed associated with diatoms and cyanobacteria through extracellular polymeric material, in some instances forming bulky structures (Fig. 3, panels A–D). Diatoms became differentiated within the external part of the microbial aggregates, protruding from the polymeric matrix. The microscopical analysis together with the identification based on morphological characteristics (Round et al., 1990; Cantoral-Uriza and Sanjurjo, 2008) enabled the recognition of the diatom species within the order Naviculales: *Navicula perotii, Navicula veneta, Navicula hodgeana, Synedra* sp.*, Amphora copulata,* and *Halamphora* sp. (Fig. 3, panels E–K).

The second layer there contained an explosion of the phylum Cyanobacteria; almost half of the reads in this area are associated with the genus *Coleofasciculus,* a well-studied and widespread marine mat. This finding is consistent with those of previous studies that identified this layer as primarily phototrophic and green in color. SEM images also confirmed the dominance of the Cyanobacteria *Coleofasciculus* sp. in the oxic zone of layers 1 and 2 (Temes-Casas and Seoane, 2000; Kim et al., 2017; Huang et al., 2020) (Fig. 3, Panels L-O). These filamentous Cyanobacteria form dense cross-linking networks joining themselves with diatoms and other prokaryotic cells to provide the backbone for the growing mat. This activity also favored the development of organosedimentary structures in the deeper zones through carbonate precipitation and biomineralization (Casal et al., 2020).

Layers 3 and 4 evidenced a decrease in Cyanobacteria and a notable increase in reads associated with Firmicutes, which were represented mainly by the anaerobic genus *Halanaerobium.* Certain *Halanaerobium* strains are known to ferment complex organic matter to produce intermediate metabolites for other trophic groups such as sulfate-reducing bacteria (Ivanova et al., 2011), whose presence has been previously predicted in these layers (Farías et al., 2013; Toneatti et al., 2017). The sulfate-reducing delta-proteobacterium *Desulfotignum* ranks fourth in read abundance in both layers. SEM observations confirmed the change in microbiome landscape wherein a lot of diatom frustules were colonized by bacteria, probably due to the increase in the concentration of nutrients as well as the availability of surface for colonization (Fig. 3, panels V–X). The SEM–energy-disperson–spectroscopy analysis correlated the high ratios of Si and O, as the main constituents of the frustules (Figs. S2 and S3).

Finally, in layers 5 and 6 the abundance of sulfate-reducing bacteria reached its maximum point, represented mainly by *Desulfotignum* and *Desulfofustis.* Archaea (Euryarchaeota) also dominated in these layers. We need to note here that in this zone crystalline structures were copious and existed in the form of aragonite (CaCO_3_) and magnesium-calcite [CaMg (CO_3_)_2_] crystals, as deduced by the abundances of the constituent elements through energy-disperson–spectroscopy mapping (carbon, calcium, oxygen, magnesium; Fig. S4). This elevated carbonate could be due to a spatial differentiation of communities as mats growing over time (Wilmeth et al., 2018). Taxa with higher motilities have the potential to move to more habitable environments when conditions become less favorable. The green phototrophic layer of STs has been found to exhibit motile behavior in reaction to light availability and ultraviolet radiation within the mats (Farías et al., 2013). Carbonate precipitation can potentially limit light availability in ST mats, forcing motile photoautotrophs to move upward, thus differentiating the organic-rich and carbonate-poor layers (layers 1–4) on top of carbonate-rich, organic-poor layers below (layers 5 and 6).

The morphotypes of bacteria observed in the images (Fig. 3, panels P–U) may be coincident with dominant bacteria found in the metagenomic analysis—*i. e., Rhodohalobacter* sp. (oxic layers), a gram-negative, rod-shaped, aerobic, and immobile bacterium (Han et al., 2018); *Psychroflexus* sp. (oxic layers), a chemoheterotrophic, and strictly aerobic gram-negative bacterium with variable morphology involving rod-shaped or coiled filaments of indeterminate length (Bowman et al., 1998); *Gracilimonas* sp. (anoxic layers), a gram-negative rodshaped, aerobic, and facultative anaerobic bacterium (Choi et al., 2015); and *Halanaerobium* sp. (anoxic layers), a strictly anaerobic, short, rod-shaped bacterium (Liang et al., 2016; *cf*. Fig. 3, Panel U).

### 2.2. Light utilization throughout the layers of the ST

In a previous publication, the light profile and distribution of photosynthetic pigments helped to differentiate zones in the mat (Toneatti et al., 2017; Fig 8, panels B, D). Chlorophyll was reported to reach a maximum in layer 1, and decreased approximately exponentially at depths >1 mm. The Cyanobacteria-specific accessory pigment phycocyanin had been previously detected mostly in layer 2. In contrast, bacteriochlorophylls (Bchls) a and c coöccurred in layer 4. Bchls a and c were generally slightly lower (but with overlap) in the layers with the locally increased chlorophyll-a content. On the basis of this consideration, the mat was divided into three main zones: an oxic, UV-stressed zone between 0 and 2 mm, where UV and blue light are intense and chlorophyll-a reached a maximum (layers 1 and 2); a transitional zone between 2 and 5 mm dominated by IR light and Bchl a and c pigments (layers 3 and 4); and finally, a low-IR zone between 5 and 7 mm (layers 5 and 6), where pigment abundances were the lowest. This last zone was also anoxic and rich in sulfidic acid. A previous report had indicated that pigments directly involved in photosynthesis were statistically overrepresented in layers 2–4 of the ST relative to the abundance in the lower layers (Farías et al., 2013).

The metagenomic survey showcased a widespread occurrence of genes for the synthesis of the UV-protecting pigments bacterioruberin, ß- and ε-carotenes, and ectoine. These pigments are concentrated in the first two ST layers and then diminish with depth (Fig 4, Panel A). The ß- and ε-carotene genes maintain a very high density in layer 1 and then decrease by half in the second layer (Table S3). According to previous reports, bacterioruberin, ß-carotenes, and ectoine could protect microorganisms from near-UV and high visible-light damage by sunscreening or by quenching triplet-state photosensitizers and reactive oxygen species (Sandmann et al., 1998; Shahmohammadi et al., 1998; Bownik and St pniewska, 2016). The concentration of these pigments, as well as those of the CPF effecting photoreactivation (Fig. 4, Panel B), was significantly higher in the first layer of the microbial mat, which depth faces continuous light exposure during the day. These UV-screening and oxidant-quenching pigments in the photic UV-stressed zone act as a UV-shield, preventing direct sunlight from adversely affecting the lower photosynthetic layers for the benefit of all members of the consortia by reducing UV-penetration and the persistence of damaging oxidants in the mat. Furthermore, that higher proportion of genes encoding the CPF and other protective pigments in the UV-stressed zone coincides with a previously described correlation between the abundance of those loci and the sunlight intensity and/or photoperiod occurring in aquatic environments (Singh et al., 2010; Alonso-Reyes et al., 2020).

**Fig. 4.**
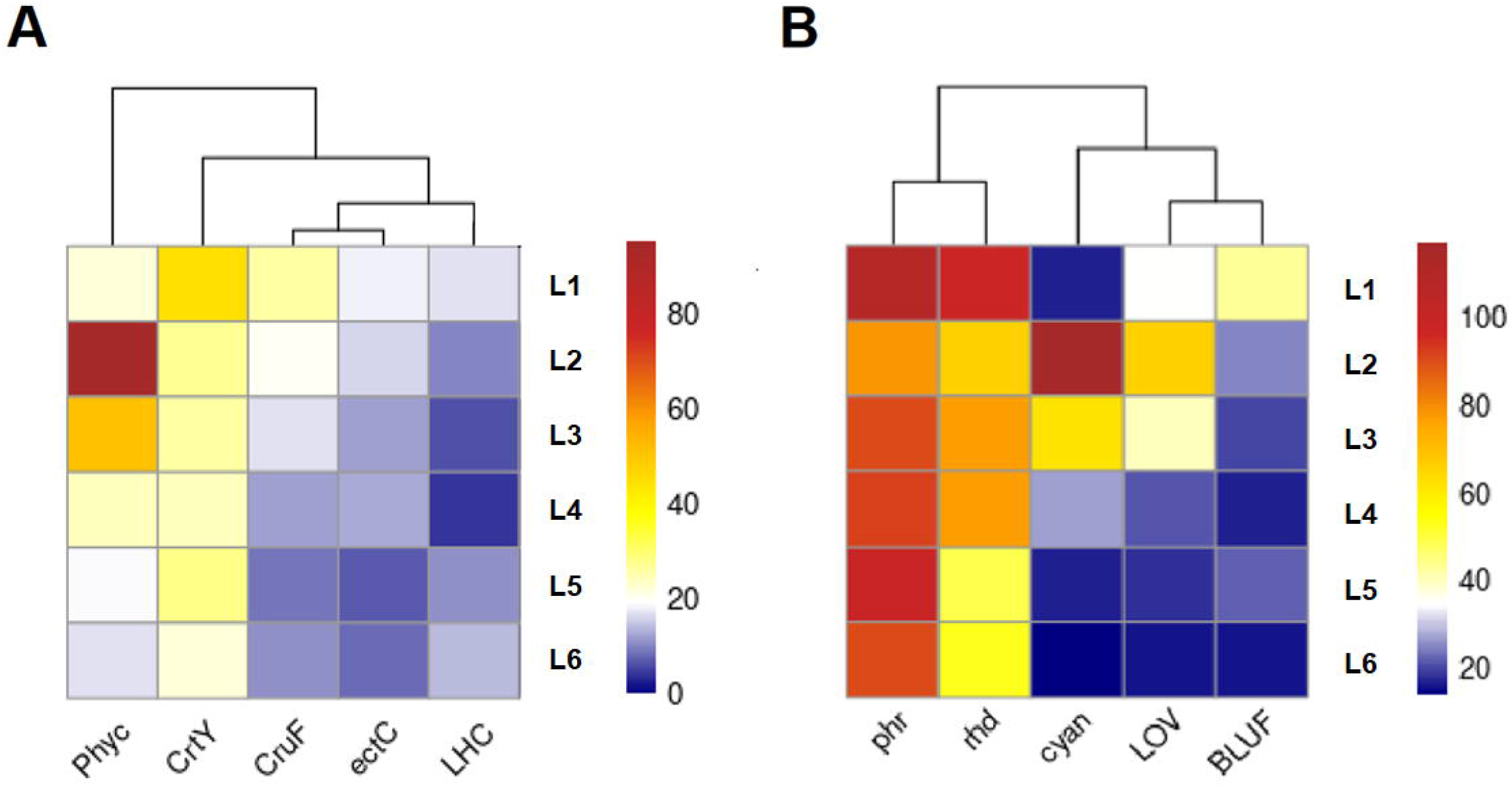
Heatmap of pigment (Panel A) and photoreceptor (Panel B) abundances. In the two panels of the figure, for each of the layers denoted on the left *ordinate*, the corresponding percent abundance of the relevant genetic locus indicated on the *abscissa* is expressed as a heat-map element, with the key to the spectrum of heat-map colors from 20% to 100% demarcated vertically to the right. The raw abundance values were recorded as normalized counts per millon of mapped reads. Phyc, phycobilisome protein; CrtY, lycopene cyclase; CruF, bisanhydrobacterioruberin hydratase; ectC, L-ectoine synthase; LCH, light-harvesting–complex protein; phr, photolyase family conferring phororesistance; rhd, microbial rhodopsin; LOV, light-oxygen-voltage–sensing domain; BLUF, blue-light-using-FAD (flavin-adenine dinucleotide) domain

To complement the previous pigment studies performed on the ST, we searched for a chlorophyll-binding domain within the light-harvesting complex (Fig. 4, Panel A) and found that that gene reached the highest abundances in layers 1 and 6. All of those sequences were associated with diatoms from the clade Bacillariophycidae (Table S3), which group in fact could be the main source of chlorophyll in layer 1 as the Cyanobacteria are poorly represented. The genes for the synthesis of phycobiliproteins—which pigments are related to the oxygen-driven photosynthesis of Cyanobacteria and red algae—exhibited concentrated abundances in layer 2, which stratum is in fact dominated by Cyanobacteria (Fig 2). Thus, the chlorophyll peak observed in layer 1 by Farías et al. (2013) was clearly due mainly to eukaryotic activity (diatom algae), whereas bacteria first start to dominate photosynthesis in layer 2.

In the absence of UV exposure, IR is the main source of photons for layers 3–6 (Farías et al., 2013), indicating that these layers are also photosynthetically active. Because of the strong competition for IR light, an effective absorption in that wavelength range coupled with the metabolic tools adapted to the hypoxic environment would constitute a selective advantage for survival and proliferation in the central layers. The shotgun metagenomics accordingly revealed that the Rhodobacteraceae was the main photosynthetic group after the Coleofaciculaceae (Cyanobacteria). In agreement, a previous publication had determined that the □-Proteobacteria from the Rhodobacteraceae family were able to carry out anoxygenic photosynthesis and thus might act as the main bacterial photosynthetic group by using near-IR light in layers 3 and 4 (Toneatti et al., 2017). Very probably most of the peak of Bchl a previously found at a 4.5-mm depth (Farías et al., 2013)—along with the Bchl-c peak from the green-sulfur-bacterium phylum Chloroflexi—was derived from these purple photosynthetic bacteria.

Accordingly, the photoreceptor profile throughout the layers was studied, including those capturing blue light (the light-oxygen-voltage–sensing domain [LOV], the blue-light-using-FAD [flavin-adenine dinucleotide] domain [BLUF], and CPF), blue and green light (the rhodopsins), red light (the phytochromes), and a varied spectrum (cyanobacteriophytochromes). With the exception of the phytochromes, all the other photoreceptors were found in the ST. The general abundances of these pigments were CPF > rhodopsins > cyanobacteriophytochromes > LOV > BLUF (Fig. 4, Panel B).

The gene for CPF (involved in UV resistance through photoreactivation) was found in a broad variety of taxa—such as the *Bacteroidetes,* Cyanobacteria, Chloroflexi, Proteobacteria, Firmicutes, unknown bacteria, and unknown taxa (Table S2). Moreover, the gene occurred at a high density in the first layer and then maintained similar abundances throughout the mat, indicating the persistence of that trait even in deep nonirradiated members of the consortia. This ubiquitousness is probably due to the ability of these microorganisms to migrate towards the surface exposed to radiation, or pure sunlight at some point in their life cycles; alternatively, that resistance trait could be thoroughly fixed in the environment of the source of those taxa. In a previous publication, we discussed the persistence of this gene family in highly irradiated regions such as the HAAL environment (Alonso-Reyes et al., 2020). We would propose here that CPF genes continually present in species are a reflection that the ecologic pressure of radiation on them from the environment under certain geographic conditions (latitude, altitude, and orography) has not changed appreciably for decades in the region sampled.

CPF sequences were classified by subfamily through the use of the sequence-similarity-network method (Gerlt et al., 2015; Figs. 5, and S5), with the abundances being considered same as those of their respective contigs. The groups *Drosophila-Arabidopsis-Synechocystis-human* (DASH) cryptochromes, iron-sulfur bacterial cryptochromes, and photolyases (FeS-BCP) had the highest abundances. A third of the sequences were not classified (28%), while the standard cyclobutane-pyridine–dimer (CPD) photolyases were the group found in a minority (9%). The DASH cryptochromes corresponded to single-stranded-DNA–specific photolyases that in many instances had secondary biologic functions, such as cryptochromes and even as double-stranded-DNA–specific photolyases (Kiontke et al., 2020). The FeS-BCPs are prokaryotic (6-4) photolyases that also often act as cryptochromes (Geisselbrecht et al., 2012; Graf et al., 2015). Both DASH cryptochromes and FeS-BCPs are secondary photolyases complementing the standard CPD photolyases that repair CPDs, the main photoproduct of the UV irradiation. The widespread presence of cryptochrome-holding subfamilies throughout the layers instead of standard CPD photolyases evidences a major relevance of the secondary biologic functions *(i. e.,* cryptochromes) as an alternative to photoreactivation in the STs. In contrast, CPD photoproducts may be less abundant in the mats, whereas singlestrand lesions or 6,4-pyrimidine-pyrimidone photoproducts should be more commonly featured. More work will be needed to clarify the function of those components.

**Fig. 5.**
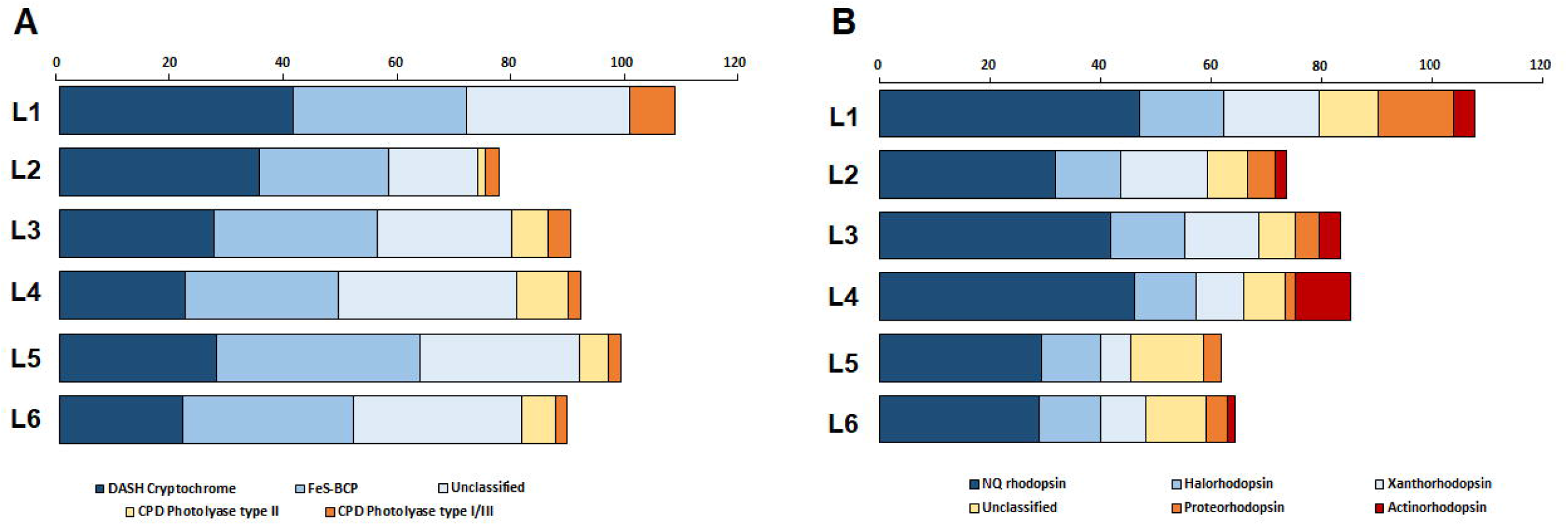
Stacked-bar chart illustrating subfamily abundance for CPF (Panel A) and microbial rhodopsins (Panel B) from the Socompa stromatolite. Each horizontal bar of the two panels, representing one of the six layers indicated on the left *ordinate*, displays the abundance, normalized as counts per millon of mapped reads and plotted on the upper *abscissa*, of each of the pigments listed below the two panels along with the corresponding color code.

The LOV-domain genes were expressed mainly in Cyanobacteria species *Coleofasciculus chthonoplastes* along with the aerobic anoxygenic phototrophic bacteria belonging to the Rhodobacteraceae family. Cyanobacteriophytochromes sequences all belonged to *Coleofasciculus chthonoplastes*, whereas the genes coding for BLUF photoreceptors were also detected mainly in the Proteobacteria and Bacteroidetes phyla and the Rhodobacteraceae family. The LOV and BLUF proteins have been associated with a great diversity of functions, including regulation of photosystem synthesis, the photoavoidance response, biofilm formation, circadian-clock regulation, light-dependent virulence, cell attachment, photoadaptation, phototactic motility, chromatic acclimation, and light-dependent cell aggregation (Metz et al., 2012; Masuda, 2013; Park and Tame, 2017; Fushimi and Narikawa, 2019). The remarkable density of sequences associated with these photoreceptors that is concentrated in the first two layers conveys an idea of the fundamental role of light in the performance of numerous biologic functions in this sector of the ST.

The highest abundance of prokaryotic rhodopsin genes was observed in the first layer, mostly from the phyla Bacteroidetes, at 62%, and Balneolaeota, at 30% (Fig. 4, Panel B, Table S2). Rhodopsins are blue-to green-light photoreceptors that can support a variety of cellular functions, including adenosine triphosphate synthesis, substrate uptake, ion pumping, sensing, and survival (Gómez-Consarnau et al., 2010, 2016; Steindler et al., 2011; Gushchin and Gordeliy, 2018). The concentration of rhodopsin genes in layer 1, situated above the bacterial-chlorophyll maximum in layer 2, may relate to the light wavelengths maximally absorbed by rhodopsins, which attenuate more rapidly than those maximally absorbed by chlorophyll a. Because this circumstance is complemented by the considerble harm to photoautotrophic bacteria caused by UV light, the phototrophic niche of layer 1 is mainly exploited by diatoms and rhodopsin-bearing photoheterotrophic bacteria capable of using light to produce enough energy to sustain a basal metabolism (Gómez-Consarnau et al., 2019). The reporting of the presence of rhodopsins is not new for the Socompa ST: a previous publication detailed the isolation and characterization of a green-tuned proteorhodopsin belonging to the *Exiguobacterium* sp. S17 strain isolated from the mat (Albarracín et al., 2016b). Likewise, Gorriti et al. (2014) detected rhodopsin genes in *Salinivibrio* sp. strains isolated from STs.

When inspected through the MicRhoDE database (Boeuf et al., 2015), all rhodopsins are classified as light-driven ion pumps that participate in energy production and are distributed among different subfamilies (Table S4). The majority of the sequences clustered in the NQ subfamily, a new group of rhodopsins with a demonstrated capability of extruding sodium and other ions (Kwon et al., 2013; Gushchin and Gordeliy, 2018). The NQ family is frequently found in hypersaline environments, even in hypersaline microbial mats, which have been reported to have an abundance of genes as high as 17% (Kwon et al., 2013). In the sodium-rich Socompa environment, employing a sodium-motive rather than a proton-motive force (bacteriorhodopsin) can be advantageous for bacteria that require sodium for their growth. Furthermore, the NQ family may be involved in the regulation of the osmotic state in the microbes of the top layers, which are closer to the hypersaline environment of the lake. Another abundant subfamily was the halorhodopsins (HRs), a group of inward-directed chloride pumps that can also translocate certain other anions, such as bromide, iodide, and nitrate (Engelhard et al., 2018; Gushchin and Gordeliy, 2018). These types of rhodopsins are also typical in halophiles where the presence of those pigments is probably related to the high chloride demands within those saline environments. The HR subfamily has been suggested to play a significant biologic role in either promoting the proton-motive force to facilitate bacteriorhodopsin-assisted energy production or maintaining a cellular-osmolarity equilibrium under the high-salinity environments found in those habitats (Chen et al., 2016). A third group of notable relevance were the xanthorhodposins (XRs), which employ a second carotenoid, salinixanthin, to act as an antenna to transfer light to the retinal proteins and extend the wavelength range of the pump (Balashov et al., 2005). XRs were present at a high abundance in the first three layers, which distribution fits the light abundance in these locations. These different groups of rhodopsins were also the main ones in other microbial mats of the region, such as those of the Tebenquiche (XR > HR > NQ) and La Brava (XR > NQ) lakes (Kurth et al., 2021).

The number of genes involved in the production of the pigments chlorophyll, ß- and ε-carotenes, bacterioruberin, and ectoine were significantly more abundant in the first layer and those for the production of phycobiliproteins in the second layer, as opposed to the lower layers (Fig. 4). A similar pattern was observed for the photoreceptors. The CPF, rhodopsin, and BLUF genes reached the highest abundances in layer 1, with the latter two decreasing in subsequent layers. Cyanobacteriochromes and the LOV domains were dominant in layer 2, but then diminished. These findings revealed a broad range of light-harvesting, light-signaling, and photoprotection strategies in the UV-stressed zone consistent with the pigment abundances previously reported. The results also demonstrate that light is a determining element in the distribution, diversity, and abundance of pigments and photoreceptors in the Socompa ST.

That the light-absorption spectra of the Socompa ST communities are well-tuned to their light environment is also notable. Photoheterotrophic bacteria carrying rhodopsins dominate the absorption in the first layer, where blue and UV light is at its maximum and thereafter rapidly declines. As a result, red light penetrates in the second layer where aerobic photoautotrophic bacteria use chlorophyll. Finally, IR light—it not having been absorbed previously—penetrates to the third and fourth layers, where anaerobic photoautotrophic bacteria with bacteriochlorophylls dominate the absorption. Thus, photoheterotrophic, aerobic photoautotrophic, and anaerobic photoautotrophic bacteria can coexist by selectively absorbing different parts of the light spectrum.

Cyanobacteria, along with associated diatoms and certain other eukaryotic algae, are the main agents of primary production in many microbial mats, via oxygenic photosynthesis (Vincent, 2002; Pepe-Ranney et al., 2012). With respect to the Socompa microbial mat, our previous pigment analysis and the present metagenomic survey revealed that bacteriochlorophyll and rhodopsins were also quantitatively essential and could overcome the participation of chlorophyll in light-harvesting. Bacteriochlorophyll and proteorhodopsins, the latter being also rhodopsins, have been recently proposed to potentially absorb at least as much light energy as does chlorophyll in certain ecosystems where the cellular energy yield by those two pigments may be sufficient to sustain bacterial basal metabolism (Gómez-Consarnau et al., 2019).

According to an ecological theory, niche differentiation over a spectrum of resources reduces competition between species, thereby promoting their coexistence (Macarthur and Levins, 1967; May and Arthur, 1972; Rueffler et al., 2006). In particular, the light spectrum is an essential selective agent for the ecology and evolution of microorganisms and is a major determinant of the species composition of communities (Stomp et al., 2007). Within that context, the coexistence of multiple light-capturing mechanisms is consistent with the selective ability of different microbes to use energy in distinct parts of the light spectrum, creating opportunities for niche differentiation. This segregation of the light spectrum is clearly observed in the Socompa ST. A pronounced attenuation of UV and blue light in layer 1 due to the combined effects of intense scattering and absorption, causing the light color to shift towards the red, and even the IR, region of the spectrum in the deeper layers. In those layers, bacterial species containing chlorophyll and bacteriochlorophyll would be capable of harvesting light at even those longer wavelengths, thus more closely corresponding to this red and IR niche.

In order to determine certain of the biologic functions associated with photoreceptors, we analyzed the effector domains of those modular proteins containing photoreceptor sequences. In this instance, only the LOV and cyanobacteriochrome domains were taken into consideration because other photoreceptors had incomplete sequences (*i. e.*, truncated contigs) or were found as unique proteins and/or domains. The LOV- and cyanobacteriochrome-effector domains were histidine kinases, methyl-accepting chemotaxis proteins (MCPs), guanyl cyclases and GDFF-EAL domains (Fig. 6, Panel A). The functions previously described for these motifs involve a two-component system, chemotaxis, biofilm formation, cyst formation, and a UV-stress response. (Kalia et al., 2013; Salah Ud-Din and Roujeinikova, 2017; Sarenko et al., 2017; Hallberg et al., 2019).

**Fig. 6.**
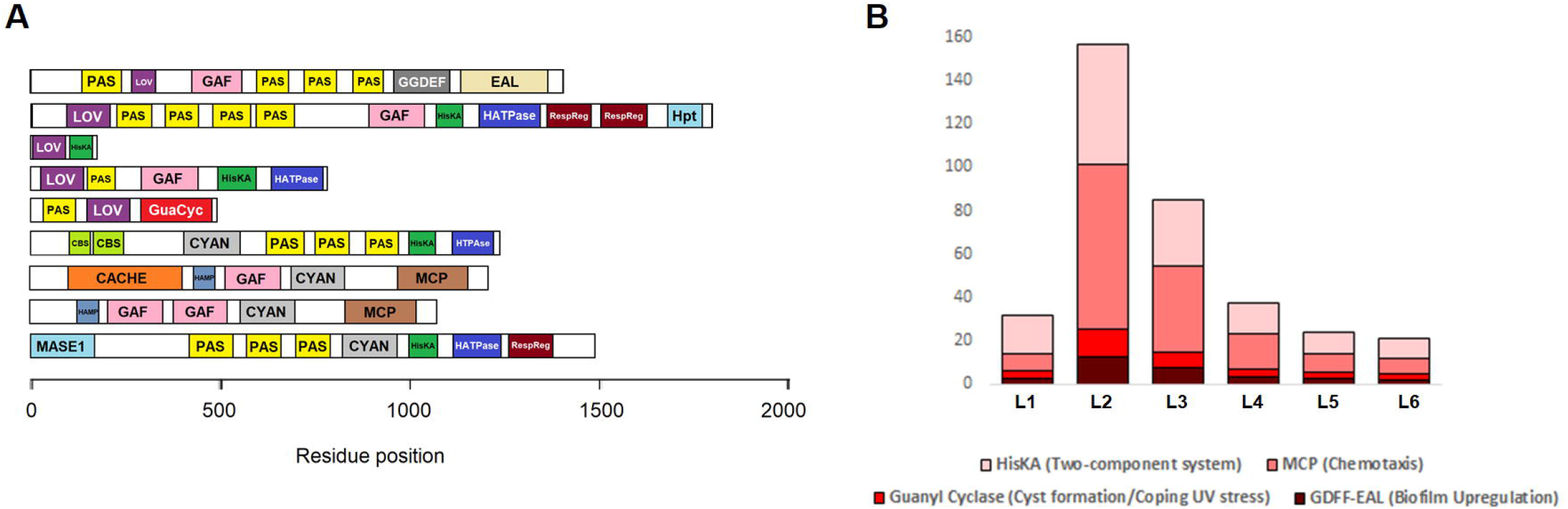
Distribution of functional domains within the Socompa stromatolites. (Panel A) Domain architecture of selected microbial photoreceptors. In the panel, the horizontal bars represent the position of the light-absoption wave lengths for the various residues present as indicated on the *abscissa* below. The widths of the component elements, whose abbreviations follow, reflect their relative abundances. The domain analysis was performed by means of the pfam service. Proportions are are reflected in the bar widths. PAS, PAS domain; LOV, LOV domain; GAF, GAF domain; HisKA, histidine kinase; HATPase, GHKL domain; RespReg, response-regulator–receiver domain; Hpt, histidine-phosphotransfer domain; GGDEF and EAL, domains named after the conserved amino-acid motifs found in their diguanylate cyclases; GuaCyc, adenylate- and guanylate-cyclase catalytic domain; HAMP, domain found in histidine kinases; CBS, CBS domain; CYAN, cyanobacteriophytochromes; MCP, methyl-accepting chemotaxis protein; CACHE, cache domain; MASE1, predicted integral-membrane sensory domain found in histidine kinases. Abundance layer by layer of photoreceptor effector domains found in the metagenome of the Socompa stromatolite. (Panel B) Abundance layer by layer of photoreceptor-effector domains. Each of the vertical bars in the panel, representing the six stromatolite layers, illustrates the fractional distribution within that layer of the four functional entities stated below the bars—namely, HisKA, the two-component system; MCP, the methyl-accepting chemotaxis protein; guanyl cyclase, an enzyme promoting cyst formation and coping with UV stress; and GDFF-EAL, a domain for biofilm upregulation. The height of each bar represents the sum of the abundances of the constituent domains in normalized units for that layer.

If we cut off every effector domain from their source sequences and sum their respective abundances separately, we can gain a notion of the functions more affected by light in every layer. For that purpose, we plotted the abundances of each of these domains in normalized units for every layer (Fig 6, Panel B). A peak is present in the second layer, where the greatest concentration of these functions occurs under the influence of light. The notable presence of chemotaxis in the second layer—which in this instance is essentially a phototaxis for having been triggered by the action of light—is consistent with the motility previously reported for the cyanobacterial layer (Farías et al., 2013).

The Socompa microbial mat constitutes a dynamic community in which microorganisms are capable of motility, thus modifying their position in the mat in search of favorable environmental conditions such as light intensity and redox potential. The color of the mat surface has been reported to change (from pinkish to green) after removal from the natural HAAL sunlight and transfer into shade (Farías et al., 2013). The resulting color changes were linked to vertical shifts in the distribution of the dominant Cyanobacteria *(Coleofasciculus* sp.) in the ST: whereas that population was concentrated in layer 2 in the sample exposed to full ambient light at 3,500 m above sea level, those bacteria became concentrated in the top layer in the shade-incubated sample. This shift probably occurred because the distribution of the dominant cyanobacteria in the STs is strongly controlled by ambient illumination, most likely through negative phototaxis away from UV-B, as our metagenomic results indicate a high concentration of methyl-accepting– chemotaxis–protein (MCP) domains associated with those of photoreception. Although the migration of Cyanobacteria deeper into the ST helps them lower their exposure to potentially harmful UV-B light, that movement nevertheless results in a lower exposure to the light that they require for photosynthesis.

### 2.3. Metabolic profile of the ST

As already stated in the previous section, the genes and pigments related to oxygenic photosynthesis were statistically overrepresented in the top two layers relative to the lower layers. As to energy metabolism, the gene profiles for aerobic and anaerobic respiration indicated that the pathway was primarily oxygenic in the first layer of the mat (Fig. 7). Indeed, as measured by microsensors, the first mat layers were oxic (Farias et al., 2013) with relatively high proportions of sequences coding for the cytochrome-c-oxidase gene, with that locus being present in even deeper mat layers. In addition, taxonomic associations of the cytochrome-c-oxidase gene coxA indicated that main groups of the ST microbial community have aerobic representatives (Fig. 7, Table S5). Nevertheless, Firmicutes, Spirochaetes, Chloroflexi, and Euryarchaeota, whose abundances increase with depth, lack aerobic members within the mat. Furthermore, genes involved in anaerobic respiration (with nitrogen and sulfur compounds as electron acceptors) were detected; and when those gene abundances were added together, they exceeded the corresponding values for aerobic respiration in layers 4–6 (Fig. 7).

**Fig. 7.**
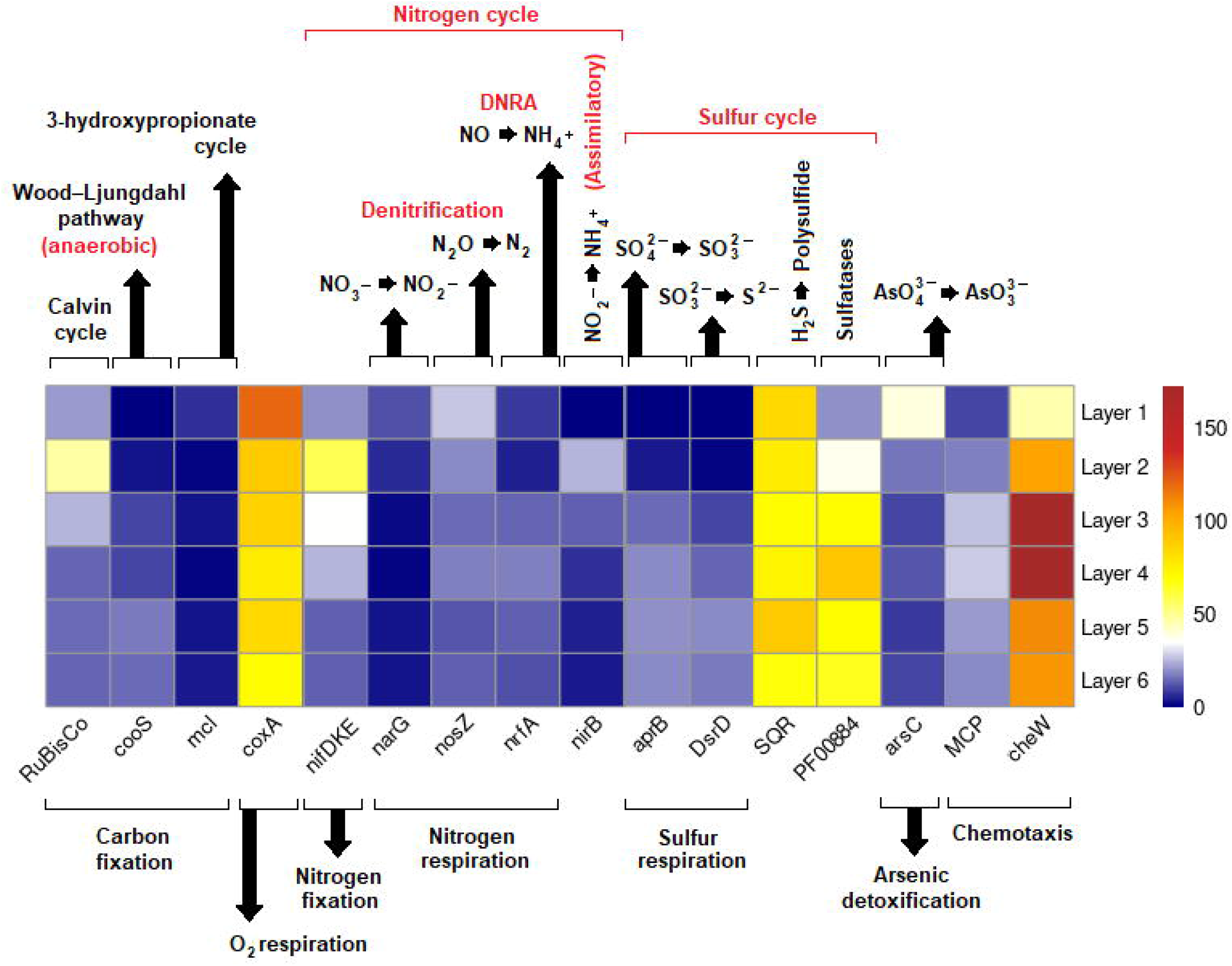
Heatmap of distribution over stromatolite depth of major functional genes in Socompa mats. In the figure, the six stromatolite layers marked on the right *ordinate* are represented vertically from the top layer down, with the corresponding heatmap color value of each respective genetic locus indicated below the *abscissa* displayed horizontally. Those genes are the major functional loci identified along with their roles in biogeochemical cycles within the different stromatolite layers of the following depths each: Layer 1, 0–0.3 mm; Layer 2, 0.3– 2 mm; Layer 3, 2–4 mm; Layer 4, 4–5 mm; Layer 5, 5–6 mm; Layer 6, 6–7 mm. The gradient from red through yellow to blue reflects the semiquantitative gene abundance throughout the six mat layers, with red representing gene overrepresentation and blue the genes with least relative abundance according to the color key to the right of the figure. The abundance range corresponding to that spectrum is reported as the normalized counts per millon mapped reads. RuBisCo, ribulose bisphosphate carboxylase; cooS, carbon monoxide dehydrogenase; mcl, malyl-CoA lyase; coxA. cytochrome c oxidase; nifDKE, nitrogenase; narG, respiratory nitrate reductase; nosZ, nitrous oxide reductase; nrfA, cytochrome c nitrite reductase; nirB, nitrite reductase (assimilatory); aprB, adenylylsulfate reductase; DsrD, sulfite reductase D (dissimilatory); SQR, sulfide-quinone reductase; arsC, ArsC family; MCP, methyl-accepting–chemotaxis protein; cheW: cheW-chemotaxis protein

Despite the relatively high presence of the coxA gene for oxygen consumption in the deep layers of the ST, the gene abundance was concentrated in the first layer (0.3 mm) and oxygen production in the UV-stressed zone (at 0–2 mm). The latter conclusion is based on microelectrode data, where the mats displayed a relatively shallow oxycline observed even during peak photosynthesis (Farías et al., 2013). In contrast, anoxic conditions below this depth are consistent with the emerging abundances of genes related to anaerobic metabolism (Fig. 7). This anoxia may be due to the lack of oxygenic photosynthesis, and/or to an even balance between oxygen production and consumption.

A range of carbon-fixation pathways were identified in all the metagenomic layers, including the Calvin-Benson Cycle (CBB), acetogenesis via the Wood– Ljungdahl (WL) pathway (reductive acetyl-CoA) as well 3-hydroxypropinate cycle (3HP). Reverse–TCA-cycle genes were scarce (data not shown). The malyl-CoA– lyase gene reached a maximum in the first layer and was among the less abundant genes, indicating the poor significance of the 3HP in the mat. the CBB was found to be more abundant in the second layer, the level where the other two CO_2_-fixation pathways had their lowest abundances. The Wood-Ljungdahl pathway gene was found to be more abundant in layers 5 and 6—those being beneath the oxic (surface) and microöxic (middle) zones—and that gene’s abundance was negatively related to those of genes enriched at the surface, while being positively related to the genes involved in anaerobic pathways (*e. g.*, aprB, DsrD).

Supplementary Table S5 summarizes the relative distribution of taxonomic groups involved in inorganic-carbon–capture pathways. The potential for CO_2_ fixation through the CBB was indicated by the gene encoding the small subunit of ribulose-1,5-bisphosphate carboxylase and/or oxygenase; these sequences were associated solely with Cyanobacteria and Bacillariophyta algae. In addition, CO_2_ fixation via the 3HP had only two sequences present in Alphaproteobacteria and Actinobacteria. Finally, the majority of the acetogens in these communities fall under the family Desulfobacteraceae and the phylum Firmicutes in a minor proportion.

The genes encoding the CBB pathway were prevalent in the metagenome, though mainly concentrated in the second layer. In contrast, the gene associated with the WL pathway was represented in the lowest layers and thus may facilitate the oxidation of CO for energy production or for coupling to CO_2_ fixation (Vavourakis et al., 2016). Heterotrophs have been proposed to utilize CO as an alternative carbon source for energy conservation in oligotrophic environments (King, 2015), and an adaptive response of that type may be necessary in the low layers of the mat. Because of the oligotrophic settings with the mats in Socompa being exposed to high UV irradiation (Albarracín et al., 2015, 2016a), a photodegradation of surface organic matter may possibly occur, thus providing CO as an alternative carbon source for energy conservation (King, 2015; Vavourakis et al., 2016). Moreover, the WL pathway is suggested to be one of the more ancient carbon-fixation channels and furthermore can be coupled to methanogenesis (Fuchs, 2011; Weiss et al., 2016). Recently, a significant contribution of the WL pathway was also reported in other HAAL microbial mats, particularly in Llamará, Tebenquiche, and La Brava; where the abundances of that pathway were similar to those of CBB (Gutiérrez-Preciado et al., 2018; Kurth et al., 2021). Even in Llamará, where layers were also studied in a separate approach, the same pattern was reported as we found in the Socompa mat: the CBB and 3HP pathways were the most abundant in upper, oxic layers of the mat, while the WL pathway was dominant in the deeper anoxic strata (Gutiérrez-Preciado et al., 2018). Since this pathway is considered the most ancestral channel for C fixation (Ragsdale and Pierce, 2008)—as evidenced by the role of the WL pathway as the major mode of carbon fixation in the modern-day HAAL microbial mats—these systems would appear to constitute even better analogues of Precambrian microbialites than other previously proposed pathways (Wong et al., 2016, 2017).

Niche differentiation with respect to CO_2_ fixation was apparent. The 3HP pathway reached a maximum in the UV-stressed first layer, where phototrophs faced struggling conditions to survive. By contrast, the WL pathway, which is for anaerobes, became concentrated in the anoxic low layers; and the CBB was dominant in the second layer over the two others, where oxygenic phototrophs accounted for the majority of photosynthesis.

As to nitrogen fixation, the data suggested that that activity was more prominent near the mat surface, with the genes for this process being negatively related to those more enriched in the deeper layers (*e. g.*, aprB, DsrD). Taxonomic assignments of the NifDKE sequences analyzed also suggested that nitrogen fixation in the STs is performed mainly by Cyanobacteria. Most of the genes were affiliated with the mat-builder *Coleofasciculus chthonoplastes* (highly abundant), while a small group belonged to a verrucomicrobia family (Puniceicoccaceae).

Nitrogenase is an ancient enzyme that remained conserved during the transition from the anoxic to the oxic biosphere (Raymond et al., 2004). Since diazotrophs may have comprised a key functional group in ecosystems on the early Earth that developed in N-depleted waters, an examination of diazotroph diversity in what is now considered as the modern analogue of the ancient STs appeared worthwhile. Prior results based on amplified nifH sequences—with that gene being the biomarker most widely used to study the ecology and evolution of nitrogen-fixing bacteria—suggested that the community with diazotrophic potential in the Socompa STs was rather diverse, comprising Delta-, Gamma- and Betaproteobacteria, Cyanobacteria (close relatives of *C. chthonoplastes*), and distant relatives of Spirochaetes and Chlorobiales (Farías et al., 2013). In the present work, we screened for NifDKE genes through shotgun metagenomics and obtained sparce results, mainly associated with *C. chthonoplastes*. This finding provides support for a key role of *C. chthonoplastes* not only in the physical construction of the mat but also in nitrogen fixation.

The genes for dissimilatory nitrate reduction to ammonium, assimilatory nitrate reduction, denitrification, and nitrogen fixation were identified in all the layers of the mat. The genes nrfA, for the former, and NirB, for the latter, were much more highly enriched in the deeper anoxic layers, whereas the denitrification enzymes nitrate reductase and nitrous oxide reductase were more abundant near the surface (Fig. 7). These findings suggest a certain partitioning of the nitrogen cycle with respect to depth. The analysis of the metagenome in the present work indicated that the Bacteroidetes, the Balneolaeota, and the Alpha-, Beta- and Deltaproteobacteria were responsible for denitrification (Table S5). In contrast, the assimilatory-nitrate-reduction gene nirB was associated with the Cyanobacteria with small traces being present in the of Gammaproteobacteria.

Genes encoding ammonia monooxygenase and hydrazine synthase were not detected. Genes already classified as unknown, or completely novel proteins, that could potentially fulfill these roles, however, could still be present in the Socompa mat; and therefore future studies are needed to investigate such a possibility. Nitrification, the microbial oxidation of ammonia to nitrite and nitrate, occurs in a wide variety of environments and plays a central role in the global nitrogen cycle. This process is presently known to be performed by two groups of bacteria (the Beta- and Gammaproteobacteria) and by diverse groups of Archaea (Kowalchuk and Stephen, 2001; Francis et al., 2005). Our results were also consistent with recent amplicon analyses of Socompa microbial mats that suggested the absence of those genes and that the diversity of the nitrifying community in the Socompa STs was rather low, having been restricted to two Betaproteobacterial groups *(Nitrosomonas* sp. and *Nitrospira* sp.; Farías et al., 2013). A similar pattern had been previously observed for HAAL mats of Llamará, where nitrification was barely detected and then only in mats from the oxic zone (Gutiérrez-Preciado et al., 2018).

With respect to sulfur-cycle genes, the abundances generally rose in the anoxic layers. Previous microelectrode analyses had indicated that H2S was detectable at depths from about 2 mm until at least 20 mm, reaching the most pronounced concentrations at depths from about 5 to 20 mm (Farías et al., 2013). In accordance with the microelectrode data, we have found that the dissimilatory sulfate-reduction genes aprB and DsrD were more highly enriched with depth (Fig. 7), and were associated mainly with the Deltaproteobacteria.

Sulfatases, which are involved in the hydrolysis of sulfated organic compounds, tracked inversely with oxygen concentration; those enzymes exhibited a twofold increase from the top layer down to the fourth layer and thereafter were uniformly overrepresented. As sulfatases can function in the presence of oxygen, the gradient is presumably a reflection of the availability of sulfated compounds in the mat. Although the concentration gradient of sulfated organic compounds in the mat is not known, they are produced by phototrophs (Kates, 1986) and are widespread in marine environments (Glöckner et al., 2003).

Genes encoding sulfide:quinone reductase were uniformly abundant through all the layers of the mat (Fig. 7), suggesting that sulfide can putatively be converted to polysulfide and subsequently assimilated, linking inorganic and organic sulfur cycles, as polysulfides react promptly with organic matter (Sforna et al., 2017).

The arsenic (As) metabolism of the bulk Socompa mat had been described in a previous report, indicating the Acr3 efflux pumps and the ArsC (thioredoxin type) arsenate reductases to be the dominant genes (Kurth et al., 2017). In the layer-by-layer present study, a high concentration of ArsC was detected in the first layer compared to the lower strata with a scant presence of the gene. ArsC catalyzes the reduction of arsenate [As (V)] to the less toxic arsenite [As (III)], which can then be extruded from the cell via a transport system such as ArsAB (Gladysheva et al., 1994). The water of Lake Socompa has manifested a strikingly high amount of As, thus leading to a marked exposure of the ST surface to the toxic metaloid. Nevertheless, previous measurements had reported that the ST contained just over half the As concentration of the external water (Farías et al., 2013). Therefore, the higher abundance of arsenate reductases in the surface layer must be contributing to the As homeostasis of the mat, by effectively acting as a biologic attenuator for subsequent layers. The metabolism of As was likewise studied in other HAAL microbial mats. In Laguna Tebenquiche, the mats reported had the highest As concentration in the upper half, a profile coinciding with the dark-green photosynthetic layer, whereas no As was detected in the top edge (Saona et al., 2021). In La Brava mats, As was also concentrated in the photosynthetic layer slightly below the surface in a pattern opposite to that of the Socompa mats, with the authors speculating that the relative distribution was due to the La-Brava mats’ arsenotrophic nature (Visscher et al., 2020). Further studies on the spatial distribution of both As and its respective metabolite will be needed to reveal the nature and significance of As cycling on HAAL mats.

Of interest to us was that the chemotaxis genes encoding MCP and the cheW-chemotaxis protein were highly abundant in the microöxic zone (layers 3 and 4). Organisms living in the microöxic zone, the boundary between the oxic and anoxic zones, could potentially accumulate substrates with high reductive potential in the anoxic zone and then move to the oxic zone to utilize this potential in oxidation (Mußmann et al., 2007). This tactic would require boundary-zone organisms to be motile and chemotactic. Indeed, our metagenomic shotgun results suggested high degree of microbial mobility to be occurrung between those layers and that such a form of mobility could be associated with energetic opportunities, as well as with light utilization and/or avoidance.

A rough measurement of the functional potential per organism can be made by estimating the average effective genome size (Raes et al., 2007). The program MicrobeCensus (Nayfach and Pollard, 2015) predicted an increased average bacterial-genome size at the border of the oxic and microoxic zones (1–2 mm depth): 5.2 Mb at the second layer versus 3.5–4.7 Mb for the rest of the mat (Fig. S6). This finding may reflect the increased functional complexity that would be needed for survival under the more adverse conditions at the former depth, as had been previously observed in the genome of a marine Beggiotoa occupying a similar niche (Mußmann et al., 2007).

## 3. CONCLUSIONS

Through shotgun metagenomics this study has delineated the functional potential of the STs of Socompa on a millimeter scale with respect to light sensing and the main metabolic pathways. The three niches previously defined by physical-chemical parameters (light, oxygen, pH) concur precisely with the functional microbiome stratification of the mat. The abundance of autotrophic (photo and chemosynthesis) and heterotrophic microbes is clearly modulated by environmental parameters—mainly light plus nutrient and mineral availability—throughout the ST layers. An enrichment of certain functions and the area of taxa was observed at the ST surface, while other enrichments occurred at the bottom. Fig. 8, Panel A presents a scheme summarizing the major functions, pathways, and gene and taxa abundances for each layer inferred from the data. The ST metagenome encodes a variety of carbon-fixation pathways, with the Calvin-Benson and Wood-Ljungdahl pathways proposed as the main mechanisms for fixing atmospheric carbon into organic carbon in the mat. The pattern of several genes (*e. g.*, those involved in oxygen consumption) reflects the microelectrode chemical data and pigment measurements performed in previous investigations (Fig. 8, Panel B). The relative abundance of anaerobic pathways (*e. g.*, sulfate reduction, denitrification) even at the surface where oxygen levels were high points to the possible existence of putative surface suboxic microniches in Socompa mats. Different niches have also been determined with regard to the use of light for energy production. Mat structure is likely to adapt to the Socompa uniquely extreme environment with genes encoding mechanisms to protect against UV irradiation and As in the first layer serving as a protective shield. In addition, the mat metagenome contains a high abundance of genes related to sulfur and nitrogen respiration in the lower layers. These characteristics coupled with the presence of the Wood-Ljungdalh pathway suggest that the ST mats are potentially modern analogues of ancient STs. Vertical microbial mobility throughout the ST may also occur as a result of physical-chemical and biologic gradients. Pigmented bacteria (*e. g.*, Bacteroidetes and Cyanobacteria) play major roles as ST builders and primary producers that sustain a diverse community of heterotrophs.

**Fig. 8.**
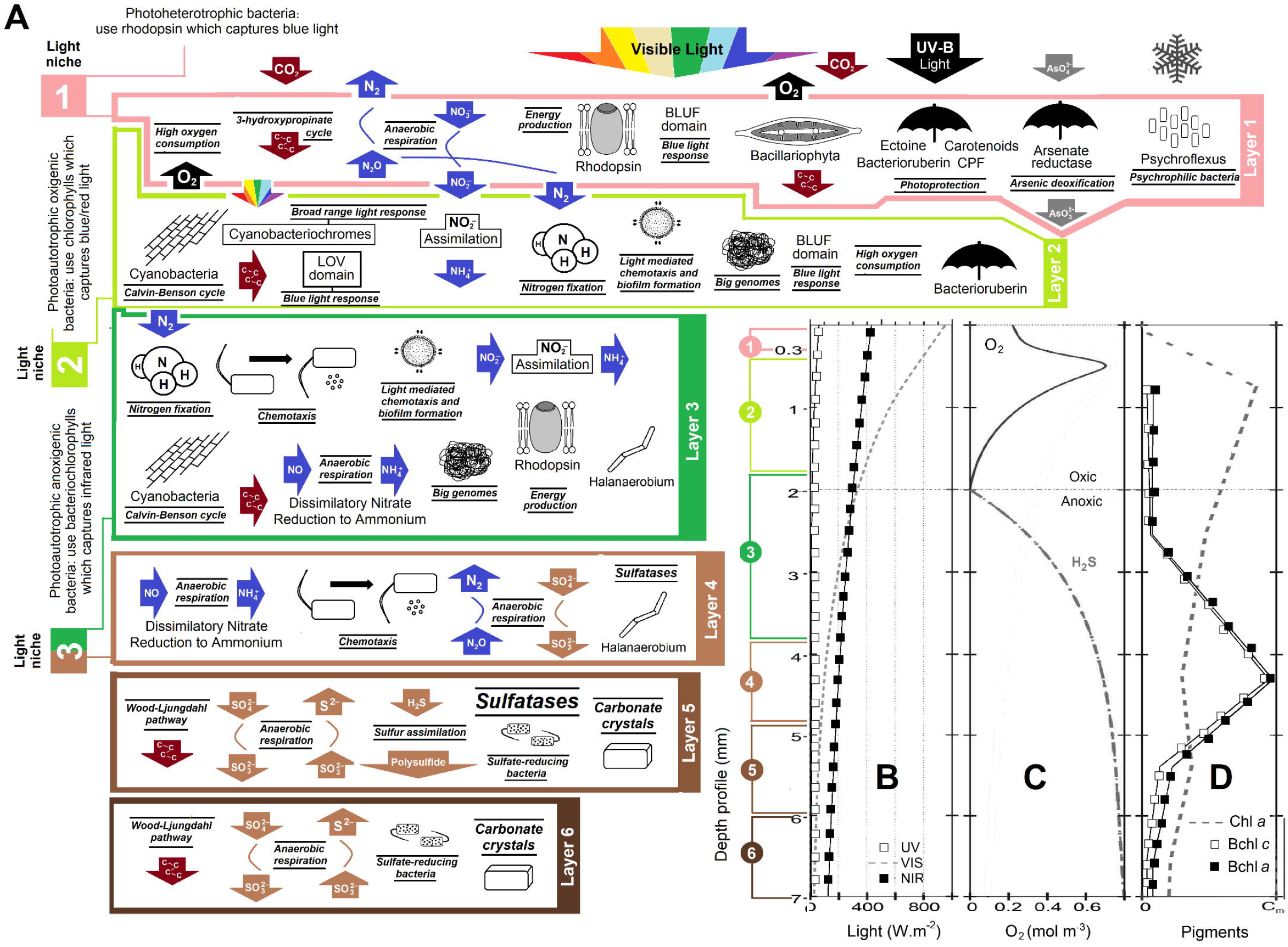
Panel A: Schema illustrating major putative microbial functions, traits, and taxa in Socompa ST as inferred from metagenomic data. To the right, six boxes representing profiles of the different depths featuring the physicochemical parameters adapted from Farías et al. (2013) and Toneatti et al. (2017). Panel B: Light profile. Panel C: Oxygen, pH, and H2S profiles. Panel D: Pigment profile. In the three panels, the parameters cited are plotted vertically with respect to the 7 ST-depth layers from top to bottom as a function of light intensity in W.m^−2^ (Panel B), O_2_ concentration in mol.m^−2^ (Panel C), and pigment levels (Panel D: dashes, chlorophyll a; white squares, bacteriochlorophyll c; black squares bacteriochlorophyll a) from 0 to C_m_ as indicated on the three respective *abscissas.*

From the results of this study, we conclude that Socompa STs are a highly sophisticated cooperative structure evolutionarily optimized for taking advantage of the resources available in spite of the constitutive extreme environmental conditions.

## 4. ACKNOWLEDGEMENTS

The authors acknowledge the generous financial support by PIUNT G603 and PICT 2019–3216 projects. FLT and DT acknowledge CAPES, CNPQ and FAPERJ. VHA, AM, and MEF are staff researchers from the National Research Council (CONICET) in Argentina. FT and DT are staff researchers from the Federal University of Rio de Janeiro. DA, SG, JMI are the recipients of doctoral fellowships from CONICET. The metagenome-sequencing project was performed in the Laboratory of Microbiology (Federal University of Rio the Janeiro, Brazil). The electron micrographs used in this study were taken at the Center for Electron Microscopy (CIME) belonging to UNT and CCT, CONICET, Tucumán. This manuscript has been released as a Pre-Print at bioRxiv. Dr. Donald F. Haggerty, a retired academic career investigator and native English speaker, edited the final version of the manuscript.

## 5. FUNDING

This study was funded by Project PICT RAICES 2019-03216 and Project PIUNT G603.

## 6. CONFLICT OF INTEREST

The authors declare that there is no conflict of interest regarding the publication of this article.

## Notes

### Competing Interest Statement

The authors have declared no competing interest.

